# Methods for broad-scale plant phenology assessments using citizen scientists’ photographs

**DOI:** 10.1101/754275

**Authors:** Vijay V. Barve, Laura Brenskelle, Daijiang Li, Brian J. Stucky, Narayani V. Barve, Maggie M. Hantak, Bryan S. McLean, Daniel J. Paluh, Jessica A. Oswald, Michael Belitz, Ryan Folk, Robert Guralnick

## Abstract

Broad-scale plant flowering phenology data has predominantly come from geographically and taxonomically restricted monitoring networks. However, platforms such as iNaturalist, where citizen scientists upload photographs and curate identifications, provide a promising new source of data. Here we develop a general set of best practices for scoring iNaturalist digital records supporting downstream re-use in phenology studies. We focus on a case study group, *Yucca*, because it has showy flowers and is well documented on iNaturalist. Additionally, drivers of *Yucca* phenology are not well-understood despite need for *Yucca* to synchronize flowering with obligate moth pollinators. Finally, evidence of anomalous flowering events have been recently reported, but the extent of those events is unknown. We use best-practices approach to annotate nearly 9,000 *Yucca* iNaturalist records, and compare the spatiotemporal coverage of this dataset with other broad-scale monitoring resources. Our findings demonstrate that iNaturalist provides unique phenology information, including delineation of extents of unusual flowering events. We also determine if unusual early flowering events impact later, typical flowering periods. Finally, we adapt a plant phenology global knowledge-store to integrate iNaturalist annotation results, supporting broadest reuse. Our approach has application to other plant groups, leveraging rapidly increasing data resources from iNaturalist to study phenology.

## Introduction

Plant phenology—the timing of plant life cycle stages such as flowering or leaf senescence—plays a critical role in terrestrial ecosystems and is known to be responsive to environmental changes (Rathcke and Lacey 1985; Ollerton and Lack 1992; Cleland et al. 2007; Chuine 2010). The fingerprint of accelerating global change, including both global-scale climatic changes and their local-scale outcomes, along with human disturbance, may show their first biotic signs in disrupted phenologies. These disruptions can have significant consequences if they lead to phenological mismatches between plants and animals that depend on them (Kudo and Ida 2013; Mayor et al. 2017).

Plant phenology data resources that cover broad scales have, until recently, only been available through monitoring networks, such as the National Phenology Network (NPN, Schwartz et al. 2012; Rosemartin et al. 2014) in the USA, which coordinates amateurs and professionals to make phenological observations. While such networks provide critical data, reporting is still sparse, since such networks often focus on key taxa or repeat sampling at a relatively small number of locations. While promising new resources are becoming available that provide wider taxonomic and spatial coverage (Silva et al. 2018), these have attendant issues with sampling protocols and with proper annotation of phenological traits and species identification.

An alternative set of resources that has yet to be broadly tapped for phenology studies comprises repositories of naturalist citizen science images. Here we focus in particular on iNaturalist^1^ as a source of phenology data because it: 1) enforces the provision of species occurrence metadata required for scientific use; 2) manages taxonomic resources, putting a premium on quality identification, and sets objective requirements for records to be considered “research grade”; 3) allows reporting of cultivation status along with annotation of traits including phenology in metadata fields, although trait annotation is still not often used; 4) is growing at a rapid and increasing pace in terms of records and species represented; 5) is directly connected with other global species occurrences aggregators such as Global Biodiversity Information Facility (GBIF), thus ensuring longer term integration and sustainability. Despite these promising attributes, best practices for use of iNaturalist and associated citizen science data resources must still be developed to realize the full value of these data streams for assessing plant phenology trends.

Here we provide a particularly salient case study, focusing on the plant genus *Yucca*, a perennial shrub or tree with distinctive flowers and with highest diversity in arid conditions in the Southwestern portion of North America, but still broadly distributed over most of continental North America. We focus on *Yucca* for three reasons. First, *Yucca* are commonly photographed and provide an excellent test case for developing proper practices for reporting needed information on phenology state. Second, *Yucca* have highly specific, co-evolved obligate pollinators and herbivores, the *Yucca* moths in the family Prodoxidae, and thus, their phenologies must synchronize to their pollinator in order to set fruit (Rafferty et al. 2015). However, the proximal environmental cues that determine *Yucca* phenology have only been examined for a few species and locations in the desert Southwest, with evidence from those studies pointing to climatic factors as determinants of the timing of inflorescence (Smith and Ludwig, 1976; Ackerman et al. 1980). Third, recent reports of anomalous flowering events in *Yucca*^2^ during Fall and Winter, and well outside of normal flowering periods (MacKay, 2013), raise the possibility that climatic changes may impact fitness, since yucca moth pollinators are presumably not yet on wing during these events. One such event was reported for Joshua Trees *(Yucca brevifolia)* in Joshua Tree National Park, but it is unknown if the spatial, temporal, and taxonomic extent was broader than these initial reports. Finally, it is unknown if those populations with anomalous events may have more restricted flowering during typical flowering phenology periods.

The first aim of our study is to derive generalized best practices for gathering phenological information from the citizen science records available from iNaturalist. As a test case, we used these general guidelines to develop a genus-specific phenology scoring rubric for *Yucca*. With this rubric, we (1) record phenological data from nearly 10,000 images of *Yucca* from iNaturalist, (2) compare these data with those available from the National Phenology Network and National Ecological Observatory Network (NEON; Elmendorf et al. 2016) in terms of spatial coverage and utility, and (3) provide an informatics workflow for integration of annotated specimens using the Plant Phenology Ontology and plantphenology.org datastore. Our second aim is to reconstruct *Yucca* phenology patterns, and to determine both which species flowered at anomalous times in autumn and well outside the normal spring or summer bloom period and the spatial and temporal extents of those anomalies and the possible impact on more typical flowering periods. We further provide a framework for downstream testing of climate factors that might determine timing of inflorescence development (i.e., flowering) in *Yucca* that can be used in future phenological investigations.

## Methods

### Data accumulation

Initially all iNaturalist records available for the genus *Yucca*, through February 15, 2019 were downloaded using function download_images from a newly developed R package, “ImageNat” (Barve 2019; The R Development Core Team 2012) that was built for this effort. There were approximately 23,600 of these records, 17,500 of which were research grade. “Research grade” records are verifiable observations where at least two participants agree about taxon identity. Verifiable observations are observations that are georeferenced, have a date of observation listed, include photos or sounds, and are not recorded as captured or planted. After exploring the number of records available per species, we decided that only species with at least 100 “research grade” records would be considered further, given our interest in ultimately developing models to predict timing of flowering; 14 species met this criterion. We also decided that species with >1,000 “research grade” records available did not need to be exhaustively sampled for the purposes of this work, and instead we randomly selected for scoring 1,000 records per species from the full set of records. All images were downloaded at the highest possible resolution available from iNaturalist in order to assure effective detection of phenology traits.

After the preliminary analysis of phenology data across 14 *Yucca* species, additional data for three species (*Y. baccata, Y. brevifolia, Y. schidigera)* were downloaded for the period of February to May 2019. These records cover the majority of the time period considered to be within the typical flowering period for these species and provided a means to examine spatial patterns of flowering phenology in the spring of 2019, after abnormal flowering periods the previous fall and winter. We restricted our taxon set for further analyses presented here to those six *Yucca* species also monitored by the National Phenology Network to facilitate direct comparisons with that data resource, shown in Table 1.

**Table 1.** Normal flowering period for the six focal species covered with overlaps between iNaturalist and National Phenology Network.

| Species name | Common name | Flowering Period | Sources |
| --- | --- | --- | --- |
| *Yucca baccata* | Banana Yucca | March-July | 1, 2 |
| *Yucca brevifolia* | Joshua Tree | March-May | 1 |
| *Yucca elata* | Soaptree Yucca | May-July | 1 |
| *Yucca filamentosa* | Common Yucca | April-August | 2 |
| *Yucca glauca* | Great Plains Yucca | May-July | 1 |
| *Yucca schidigera* | Mojave Yucca | March-May | 1 |

1 - https://www.feis-crs.org/feis/ with searches for specific species and collation from seasonal development descriptions.
2 - https://www.wildflower.org/plants/ with search for specific species and collation from “bloom information” field

### Developing best practices for gathering phenological data from iNaturalist photographs

Two authors (V. Barve and R. Guralnick) initially examined the *Yucca* photographs from iNaturalist and developed a rubric for scoring these images, focusing on flower and whole plant presence (Box 1). Definition of traits for this rubric was based on work from Brenskelle et al. (2019) and Stucky et al. (2018), who have developed a set of formal definitions in the Plant Phenology Ontology. In particular, we scored flowers, open flowers and whole plants as present or absent. We added “uncertain” as a scoring category, because initial examination revealed cases where image quality was poor or the state otherwise difficult to observe.

#### Box 1. A best practices checklist for developing a phenology scoring rubric for online citizen science photographs.

We list a series of recommendations that forms a core set of best practices when developing and implementing a scoring rubric for annotating plant phenology from online citizen science photographs. The list should serve for any online natural history photographic resource but was developed with a focus on iNaturalist.

##### Develop a consensus and standards-based scoring system

1. We strongly suggest aligning definitions and terminology to well-defined standards such as the Plant Phenology Ontology (Stucky et al. 2018).
2. We recommend utilizing scoring protocols, such as those defined by Yost et al. (2018), which were developed for digitized herbarium sheets but apply here as well. As with #1 above, those protocols ensure that scoring uses standard terms.
3. iNaturalist records often can be used to report absence, which is particularly useful for modeling climatic drivers of phenology. Definitive absence requires scoring if a whole plant is visible in the photograph, which we strongly recommend capturing.
4. We advocate a muti-scorer approach (i.e., where each image is independently scored at least twice), especially in cases where expertise is lower. In cases of conflict, an expert panel can be used to resolve issues.

##### Develop iterative training approaches and record flagging mechanisms for volunteers

1. We suggest an iterative procedure for testing volunteer’s internal definitions. This iterative procedure involves multiple testing rounds and checks for scoring consistency.
2. Consistency checks can reveal photographs that are particularly difficult to score, and discussing those photographs as a group will help refine scoring rubrics.
3. Adding a way to flag challenging records during the actual scoring assurs the scorers can bring those records to the group for discussion.

##### Develop efficient scoring mechanisms

1. Our efficiency increased markedly when we properly used dependency chains when scoring phenology, and we strongly recommend developing these. For example, if a flower was present, we did not need to score whole plant presence traits.
2. The use of custom software for image annotation proved to be a significant time-saving device, rather than laboriously shifting between an image viewer and spreadsheet. It also decreased errors in transcribing.

##### Publish data and metadata in established repositories

1. We encourage immediate publication of validated phenology annotations into repositories, such as the Global Plant Phenology Portal (plantphenology.org), to assure best-possible integration and reuse. More work to set up licensing and rights, etc., along with full metadata about methods will further improve downstream re-use.

We recruited nine additional graduate students or postdoctoral fellows (besides V. Barve and Guralnick) as part of a semester-long project at the Florida Museum of Natural History (FLMNH), most of whom were not familiar with *Yucca* before beginning the project. After a first presentation of the scoring rubric, 11 scorers were given the same 100 iNaturalist images to score as a test for consistency. We reconvened to discuss photographs where among-scorer conflict existed, and reached a group consensus of how these cases were to be scored. After a series of refinements based on inter-scorer comparisons, the 11 scorers were each given a new, identical set of 100 more records to test consistency again. At this point, scoring was greater than 90% consistent among scorers across species and traits. Finally, after these practice sets, each scorer was assigned two or three sets of 1,000 images to score on their own. The end result is that each image was scored by three independent volunteers, which is the minimum number required to both determine any potential scorer conflict and generate a majority rule assessment.

Initial scoring work was done on spreadsheets, but to hasten and help automate steps in the scoring process, one of us (Stucky) developed a software tool called ImageAnt (https://gitlab.com/stuckyb/imageant) to help with the task, which will be described more formally in a separate contribution. ImageAnt uses a simple language so that users can define transcription targets, and an order of scoring. To help make image annotation as efficient as possible, ImageAnt can use the answers to high-level questions to decide which lower-level questions to display. For example, if the user indicates that flowers are present on image, ImageAnt will not ask whether the photograph is of a whole or portion of a plant. Whether a photograph shows a whole plant or portion of a plant is only relevant in cases where flowers are absent. The ImageAnt software then saves annotations as a CSV file.

During the scoring effort, we continued to identify difficult cases and addressed those issues through iterative refinements. In particular, for the full scoring effort, we added a new scoring category in ImageAnt that enabled scorers to flag records they found particularly problematic, allowing us to go back and review these records as a group. These flagged records were different from those recorded as “uncertain”; in many cases annotators were certain in their uncertainty.

Some important decisions were made prior to the classification process that affected how images were scored and future applications of the dataset. For example, we decided not to choose a single focal plant in an image because, in some cases, there are other plants *of the same species* in the background but with a different phenological stage. To be clear, we only scored the target species (that is, the species in the photograph matching species metadata), and did not consider other *Yucca* species even if co-occuring in the same digital photograph voucher. If at least one whole plant of the target species was in view in an image, the image was scored as “whole plant present”. Similarly, we scored flowers and open flowers present if any plant of the target species (whole or partial) in the image had those states present. A supplement document (S1 - Best Practices Guide for Scoring *Yucca* Images on iNaturalist) provides full details on scoring practices.

In sum, three independent scores were captured for each of the 8,575 total images. After a further vetting to remove a subset of images that could not be scored (discussed in Results), we compiled a scored annotation dataset for each photograph. Next, in cases of conflicting reports of presence or absence of a trait, a smaller group (V Barve, Guralnick and Brenskelle) individually reviewed those inconsistencies case-by-case. After that review, the smaller group reached a consensus decision among themselves, and finalized a score for those records. V. Barve, Guralnick and Brenskelle also individually scored phenology for a spring 2019 subset of records (discussed above), and then collectively reconciled results among themselves to derive a final vetted annotation. Finally, we abstracted a set of general best practices (Box 1) usable for any phenology scoring project that emerged from our efforts.

### Comparing iNaturalist, NEON and National Phenology Network phenologies

We downloaded phenology observation data from the National Phenology Network^3^ and NEON^4^ for all *Yucca* species from both source datasets and from the Global Plant Phenology Portal^5^, which provides a harmonized set of reporting. Table 2 provides a summary of the number of observations of whole plants, or annotations from photographed plants, for each source. We also plotted spatial distribution of phenology reporting for all sources, along with temporal trends in datasets from 2014-2019 for overlapping species (Figures 1–2). We especially examined spatial and temporal patterns of anomalous blooms in fall and winter of 2018-2019 for those species with data from multiple phenology observation resources. We defined anomalous flowering as flowering well outside known bloom periods; Table 1 provides a summary of normal flowering period, along with citation source, for the key 6 species (noted above) with overlapping data in all repositories. Anything found well outside of those ranges of time were considered anomalous. As a clear example, there were multiple sightings of *Yucca brevifolia* in flower in November of 2018, which is well outside its typical bloom period in March–May. In order to visualize whether those areas with anomalous bloom periods also had *Yucca* flowering events during the typical typical blooming period, we plotted spatial patterns of blooming constrained by different time periods (Figure 2).

**Figure 1.**
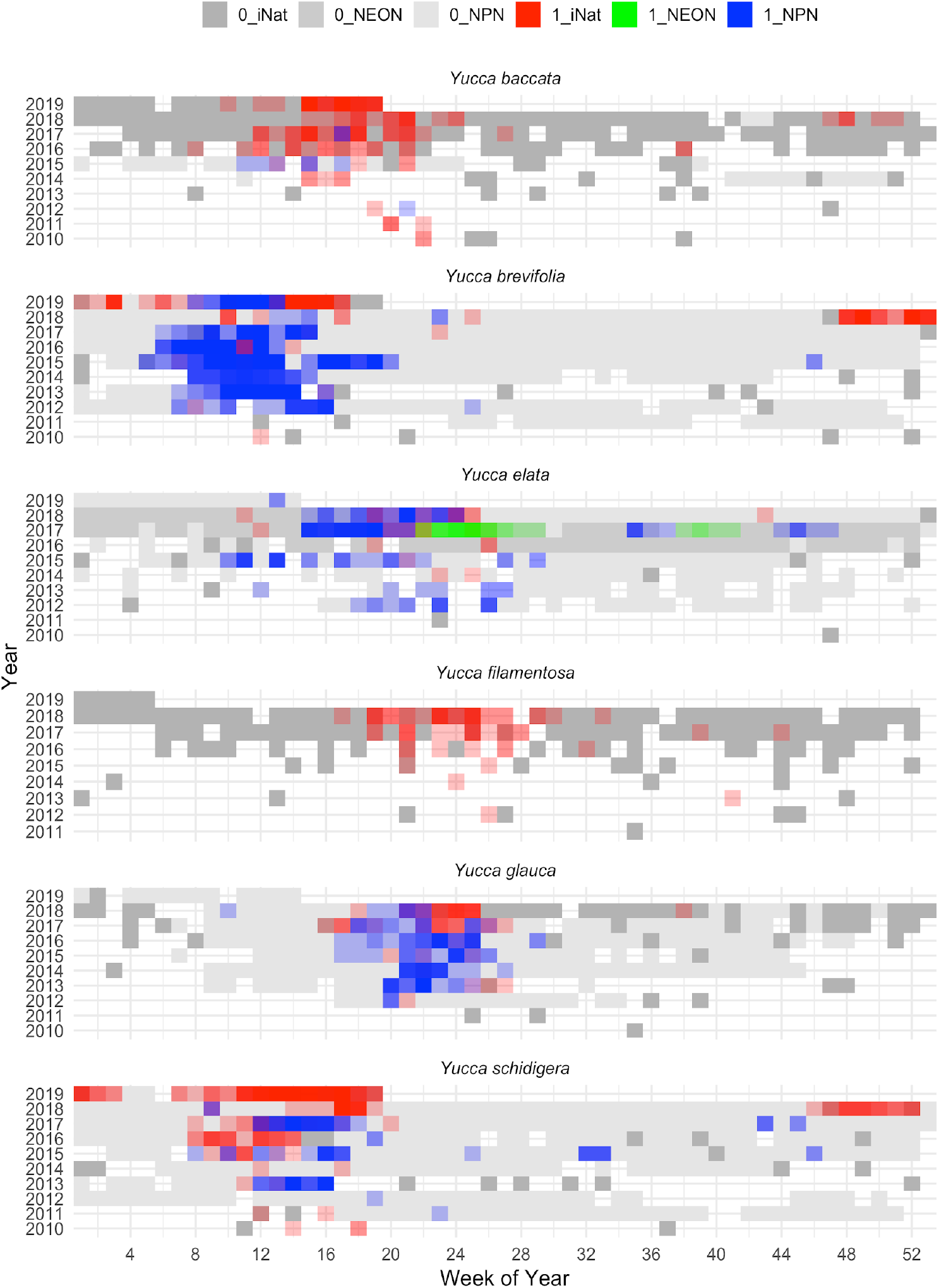
The flowering of the six focal *Yucca* species are shown here with colored boxes for each week of the year from 2010-2019. Flowering presences are indicated with different colors documented by iNaturalist (red), NEON (green), and NPN (blue). The coloring intensity indicates the number of reports from a given source during a specific time period. The colors are mixed when there are flowering presences reported from multiple sources for a single week.

**Figure 2.**
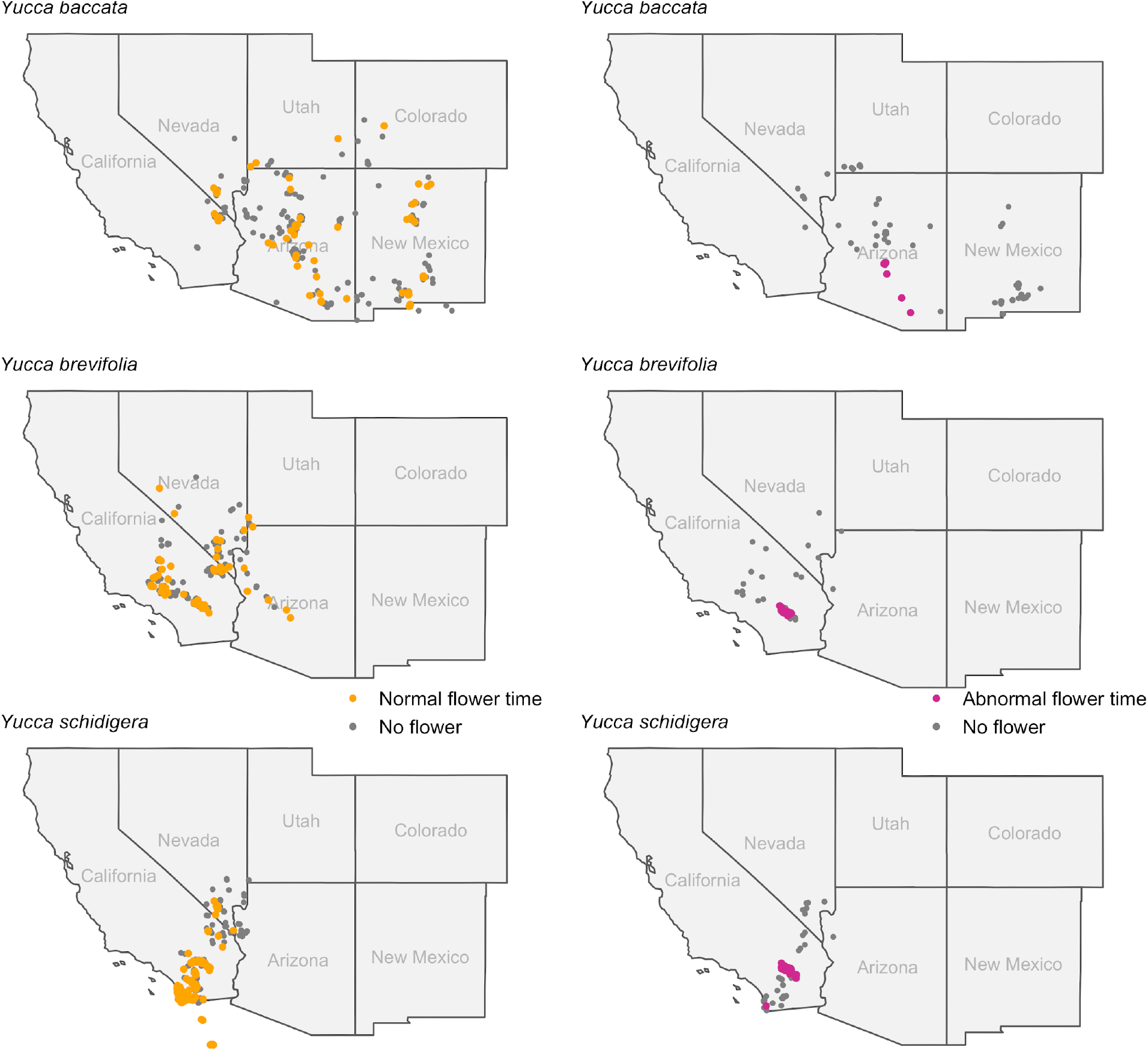
The spatial distribution of both normal flowering time (yellow dots) in spring 2019, defined broadly here to capture potential early onsets as February 11-May 2019, and anomalous flowering times (magenta dots) for the three *Yucca* species during late fall and early winter 2018-2019. Anomalous flowering events here are those occurring November 1, 2018- February 10 2019), well outside normal known time frames. These are superimposed on absences also drawn from the same time periods. Points outside of map boundary e.g. for *Y. schidigera* are located in Mexico, which is not shown here.

**Table 2.** Number data points for each focal species that had two or more sources.

| Species name | iNaturalist | NPN | NEON |
| --- | --- | --- | --- |
| *Yucca baccata* | 902 | 72 | 0 |
| *Yucca brevifolia* | 1213 | 17367 | 0 |
| *Yucca elata* | 2845 | 3730 | 3499 |
| *Yucca filamentosa* | 417 | 0 | 0 |
| *Yucca glauca* | 384 | 15478 | 0 |
| *Yucca schidigera* | 1222 | 3533 | 0 |

### Publishing iNaturalist phenology annotations

In order to make our *Yucca* phenology scores available in a reusable format, they were uploaded to the Global Plant Phenology Data Portal^5^. In preparation for ingestion, these data were converted into a spreadsheet with associated iNaturalist URIs, observation metadata such as the location and date of the observation, and descriptions of phenology using terminology from the Plant Phenology Ontology (PPO; Stucky et al. 2018). In this first round of provisioning data to plantphenology.org, we did not provide scoring reconciliation metadata. That is, we did not provide metadata indicating scoring conflict in records or how these were resolved. However, such metadata are important to report; in future iterations we plan to provide information about the scoring process. Once the data were reformatted for ingestion, they were uploaded to the portal using the ingestion pipeline developed for the PPO, described further in its GitHub repository^6^.

## Results

### Developing a scoring rubric and outcome of *Yucca* scoring

We developed a general approach to scoring iNaturalist photographs summarized in Box 1. That approach includes a set of best practices that we argue should be followed, if the intention is to create research-quality phenology data from online photographic resources. We implemented this approach, leveraging effort from eleven trained volunteers (all included as authors) who each scored between 2000-3000 images for all three traits of interest (whole plant presence, flower presence and open flower presence). All 8,575 photographs were annotated by at least two people, and most were scored by three. After initial scoring, a subset that could not be scored at all were removed, leaving a total of 8,129 that were assembled into a final dataset that captured phenology reporting for 14 *Yucca* species. We focus hereafter on the six species that overlap with reporting from the National Phenology Network.

Figure 3 shows results for scorer consistency, both per species and overall, after initial practice rounds and using the fully developed rubric. We expected that whole plant assessment would be easier for *Yucca* species that are more branched and tree-like, such as *Y. brevifolia* and *Y. elata*. However, those larger species are sometimes photographed from a farther distance, making it challenging to assess flowering phenology. Overall, our results show that scores of flowering phenology as uncertain was indeed highest in the largest species, *Y. brevifolia* (30-40%), and the second largest species, *Y. elata* also had moderately frequent uncertain scores (10%). As expected, we found the greatest conflict among scorers in detecting whole plants versus partial plants, since this can be challenging in photographs where the ground is not visible due to dense undergrowth or the angle of the photograph, or where clonality makes it unclear whether a whole individual was captured, all of which happened frequentl*Y. Yucca filamentosa* was the most challenging to score as whole versus partial plant based on its unusually high inter-scorer conflict (>15%), likely because it is located in denser, more mesic habitats in the Southeast with greater plant crowding. In sum, despite species variance, the overall rate of uncertain scoring was still relatively low, below 1-2% in a majority of species. Inter-scorer conflict was also relatively low for flowering traits, which provides evidence that the scoring rubric could be used successfully in most cases, and that the primary conflict was in assessing whether a complete individual was captured rather than recording the phenology states in the photograph.

**Figure 3.**
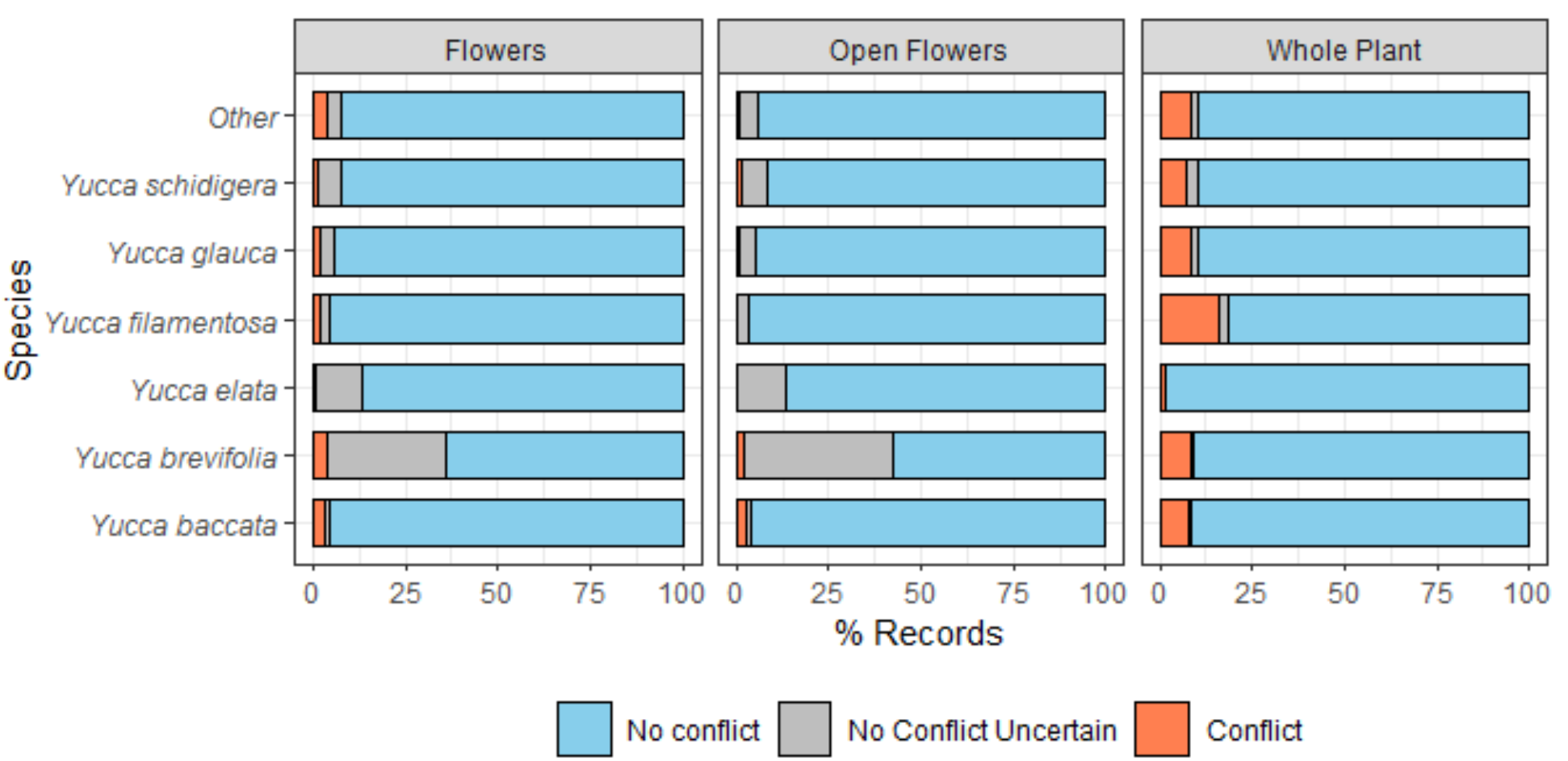
Percentage of non-conflicting, conflicting and uncertain records per species. Larger, tree-form *Yucca* often have increased rates of uncertainty in documenting flowers, while some smaller shrub species, such as *Y. filamentosa*, proved challenging for documenting whole versus portion of a plant.

Table 3 shows per-species percentages of whole plants recorded for those photographs where flowers are absent. We were particularly interested in whether photographers generally try taking photos of whole plants, since these are the ones most useful for demonstrating absence of flowers. Indeed, the vast majority of photos (>90%) are of whole plants, with no discernable bias for more or less whole plant reporting in larger, branched species and smaller, stemless shrub species. We also expected a relatively high percentage of photographs with plants in flower, given known observation bias toward recording flowering individuals. Excluding *Y. elata*, we found that the percentage of the target *Yucca* species in flower on iNaturalist ranged from 16%–26%. In the case of *Y. elata*, one iNaturalist photographer was extremely active in photographing that species whether in bloom or not, resulting in a much higher rate of recorded flower absence. Taken together, our results suggest a general bias toward observers photographing plants in flower.

**Table 3.** Percentage of whole plant photographs for those records with flowers absent and overall percentage of photographs with flowers (whether opened or unopened). Most photographers are capturing whole plants, and are biased toward those plants with flowers, as discussed in the text.

| Species | Percentage of Whole Plant photographs | Percentage of Flower Photographs |
| --- | --- | --- |
| *Yucca baccata* | 0.93 | 0.20 |
| *Yucca brevifolia* | 0.90 | 0.23 |
| *Yucca elata* | 0.98 | 0.06 |
| *Yucca filamentosa* | 0.93 | 0.17 |
| *Yucca glauca* | 0.95 | 0.16 |
| *Yucca schidigera* | 0.90 | 0.26 |

### Comparison of different observation network reporting of Yucca phenology

Figure 4 shows the spatial coverage of records from different phenology sources. Two patterns are immediately visible. First, iNaturalist records provide significantly more spatial coverage of phenology than NPN or NEON. That spatial coverage comes at the expense of repeat temporal coverage that is lacking in iNaturalist and a critical strength of NPN. For each NPN site, there are often hundreds of repeat measurements of the same individual or populations. However, the National Phenology Network also includes many repeat-sampled populations that are outside the native ranges of the species it samples, likely representing either cultivated specimens or misidentifications. This is most clear for *Y. schidigera* and *Y. glauca*. In *Y. schidigera*, the majority of sites where observations occurring in the core part of the range, but there are two sites - one in Northern Colorado and one in the Bay Area of California, that are well-outside known distributions and must be cultivated specimens. In *Y. glauca*, the Great Plains yucca, multiple reports in Southern California as well as Tennessee and south-central Arizona are all well outside the known range of the species (Althoff, 2016). Unfortunately, reporting about whether these are cultivated or wild populations cannot be easily determined in the National Phenology Network datasets.

**Figure 4.**
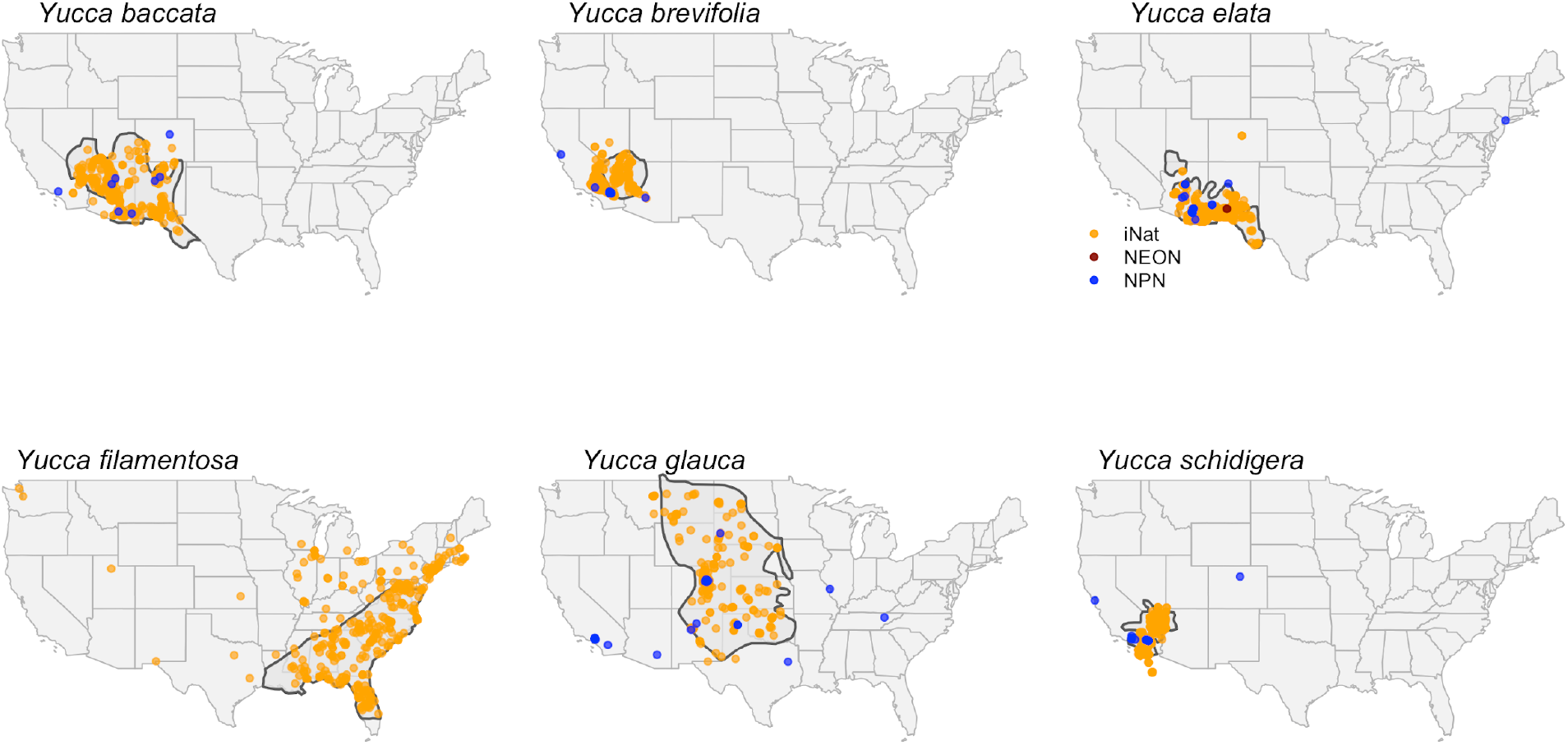
The spatial distribution of occurrences of the six focal *Yucca* species is shown with different colored dots indicating the occurrence source. Species’ range maps from eFlora.org were digitized and included in the background.

### Documenting anomalous flowering in space and time

*Y. brevifolia* and *Y. schidigera* both show a strong signal of blooming in mid-November 2018 that continued through January 2019 (Figure 1), which is well outside normal bloom timing (Table 1). Anomalous flowering is also seen in *Y. baccata*, the banana yucca, for a shorter duration during November and December of 2018. When the anomalous bloom is mapped spatially, it is striking that anomalous blooming events were spatially restricted in all three species. For *Y. brevifolia* and *Y. schidigera* the anomalous flowering is restricted to areas in and around Joshua Tree National Park, with the exception of two *Y. schidigera* with flower buds observed in very late January in San Diego County (Figure 4). *Yucca baccata*, the range of which does not overlap with the previous two species, also experienced anomalous blooming evnets during fall-winter 2018 in south-central Arizona, well outside the normal period of March-July.

### Publishing scoring results to plantphenology.org

In order to make iNaturalist-derived phenology annotations openly available, the plantphenology.org portal was reconfigured to add iNaturalist as a source for phenology annotations. We labeled the source as “Image Scoring Records from iNaturalist” and results returned are provided as either a map or table, all available for download, from plantphenology.org. The records are also available via the R package “rppo”. In all cases, the individual record results always point back to a URL for the observation record, including the photograph from which the annotation was made, in iNaturalist. Figure 5 provides a screenshot showing an example of a result return for all 14 scored *Yucca* species with open flowers present, showing mapped results.

**Figure 5.**
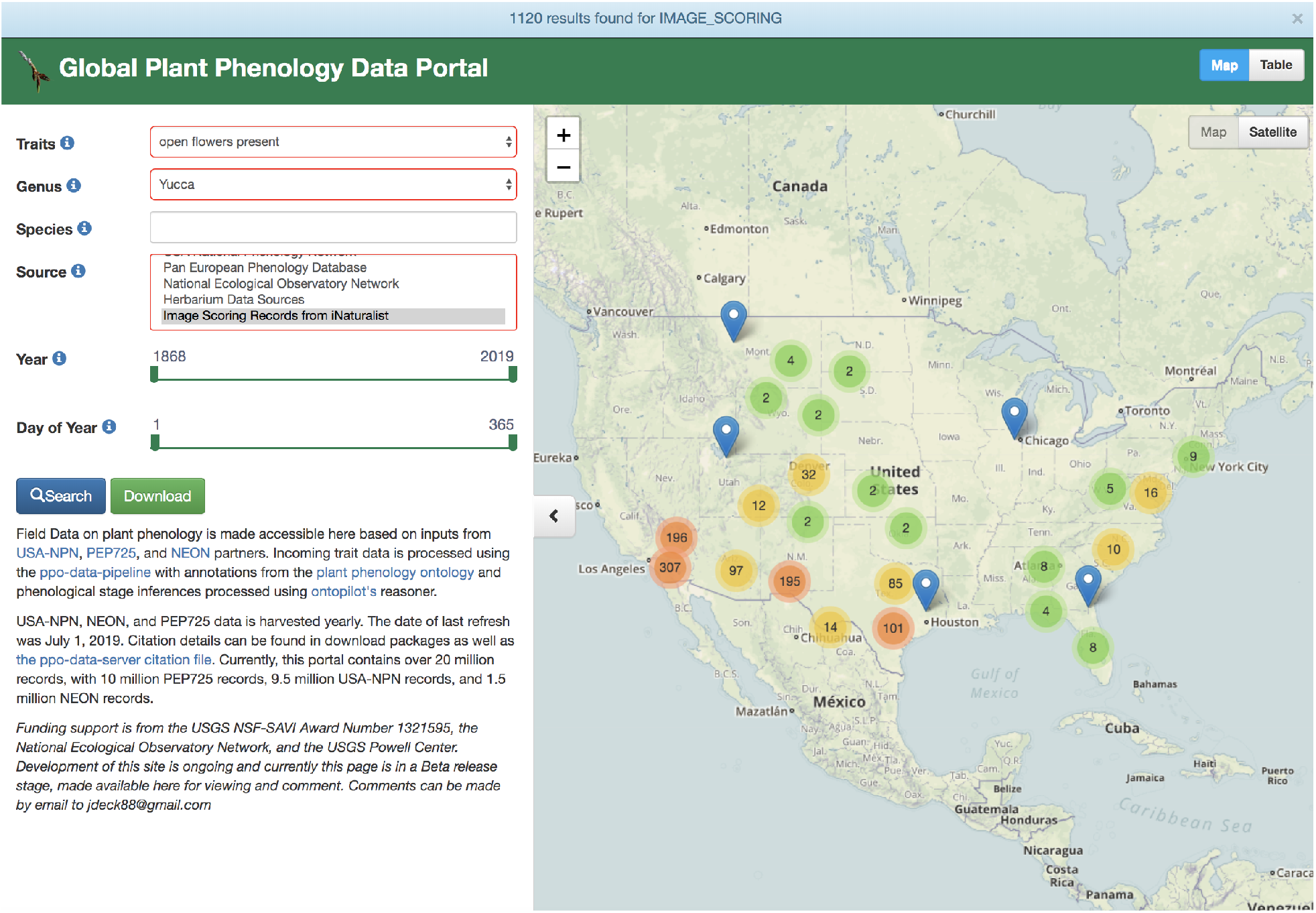
The records this project added to the Global Plant Phenology Data Portal are searchable by choosing the “Image Scoring Records from iNaturalist” source and searching for the *Yucca* genus. The portal returns a map interface by default but records can be viewed in table-mode and downloaded.

## Discussion

### The need for phenology scoring best practices from citizen science photographs

We provide a brief best practices summary for developing plant phenology scoring methods for online, citizen science photography sharing platforms in Box 1. We also provide a Yucca- specific rubric in Supplement 1. As Box 1 discusses in more detail, key practices including aligning scoring to well-defined terms in ontologies, developing an iterative scoring process, working collaboratively with volunteer scorers, and using annotation tools to increase efficiency. These best practices are essential, because scoring whole versus portions of plants and flowering traits proved surprisingly challenging for *Yucca*, based on not only our quantitative results of inter-scorer comparisons, but also from the sometimes energetic discussions that arose. We also note the importance of being able to report uncertainty, especially in tree-like species, where photographing the whole plant often resulted in reduced resolution and associated difficulty identifying flowering stage. Finally, we note that iNaturalist itself has reporting mechanisms for plant phenology, so a longer term goal is to assure that scoring efforts work in both directions, such that annotations can ultimately be fed back to iNaturalist, and to encourage further effort to denote phenology by the more than 500,000 citizen scientists who contribute to that platform.

We doubt that the scoring challenges discussed above are unique to *Yucca*, necessitating broad best practices, but developing a more specific guide to *Yucca* scoring (Supplement 1) provided some needed specificity for taxon-specific scoring challenges. As an example, our iterative, consensus-based approach led to recognition that scoring of whole versus a portion of a plant was not independent of flowering scoring. For example, it was necessary to determine whether or not the flowering stalk was living or dead in order to score whole plant presence. If the stalk was dead, but not completely in view, we scored whole plant present. Conversely, if the stalk was alive but not fully visible, we scored whole plant absent.

We also quickly determined that scoring whole versus part of plant cannot be done independent of species-level taxonomic knowledge since expected differences in growth form (e.g., clonality, caulescent habit) are needed to assess the presence of an entire individual. This proved especially important in cases where it is difficult to determine the plant’s position relative to the ground, which can happen in photographs with multiple species or of landscapes. Marginally less difficult was scoring buds versus open flowers. For that rubric, we only scored unopened flowers present in cases where a bud was clearly visible. In many cases, a stalk had formed and bracts were visible but not buds; these were scored as flowers absent. Transitional states between unopened and opened flowers and between opened flowers and senesced flowers (here scored as flowers absent) are always challenging, and more details about our rubric for those states are found in Supplement 1.

### The unique value of iNaturalist for providing spatial flowering phenology coverage

A guiding question that motivated our research was whether the coverage of iNaturalist could improve understanding of flowering phenology pattern and process compared to what is available from the National Phenology Network (NPN) and NEON, which are the key monitoring datasets usable for generating phenoclimatic models, especially in the United States. Previous work examining flowering phenology in *Yucca* has been limited to just a few sites and years (Smith and Ludwig, 1978; Ackerman et al. 1980), and no explicit models have been developed to determine if climate factors, photoperiod, or the interaction between the two can be predictive of flowering time. This question is particularly important, because if climatic factors do control *Yucca* flowering phenology timing, it is possible that climate change could create mismatches between *Yucca* and their obligate moth pollinators (Rafferty et al., 2015).

A key finding when comparing iNaturalist records to National Phenology Network and NEON monitoring is the much broader spatial extent of records found in iNaturalist. iNaturalist records may provide a good example of the power of observations collected over a gradient, as opposed to those collected via repeat sampling. Detecting non-linear trends may be greatly improved with such broad-scale sampling, as opposed to more replication across fewer sites (Kreyling et al., 2018). Additionally, we found that iNaturalist records are almost all within the core known range of species, which is not the case with the more sparsely spatially sampled NPN records. Finally, NPN records for *Yucca* sometimes appear to be cultivated specimens, but limited reporting makes detection of these cultivated specimens challenging.

Similar issues with cultivated specimens also occur on iNaturalist, but it is simple to tag records as cultivated, and those listed as such cannot become “research grade”. While we have not undertaken a quantification of how many clearly cultivated or planted specimens are not labeled as such and become “research grade”, a cursory examination of “research grade” *Yucca* observations suggests that the vast majority are indeed non-cultivated. Further, because spatial sampling in iNaturalist is much larger than temporal sampling, the problem with cultivated specimens in relation to phenology patterns may be less acute. For example, the National Phenology Network reports of an unusual flowering period for a specimen or specimens of *Y. elata* in the Joseph Wood Krutch Garden on the University of Arizona campus in the fall of 2017. This site had multiple reported days of flowering over two periods, in late August/early September, and again over multiple days in November, and is clearly visible on the *Y. elata* plot in Figure 1. No other sites for *Y. elata* recorded this unusual bloom pattern. Cultivated specimens, especially in managed gardens such as the one on the University of Arizona campus, may experience different conditions (e.g. watering regimes) from non-managed plants, desynchronizing normal phenology processes (Buyantuye and Wu, 2012). Unless information about those management regimes are known and can be included in models, such records may ultimately obscure understanding of drivers of flowering phenology. Detection of natural flowering anomalies from those occurring due to cultivation is both critical and challenging, especially in cases such as data from NPN, given lack of reporting methods and photographs usable for judging surrounding landscape.

### Documentation of restricted anomalous flowering in Yucca

Our work shows clear evidence of anomalous flowering in fall 2018 for three *Yucca* species. In all cases, that anomalous flowering is spatially restricted in extent, occurring in Joshua Tree National Park for *Y. brevifolia* and *Y. schidigera*, and in south-central Arizona for *Y. baccata*. In the case of anomalous flowering in and around Joshua Tree National Park, the only two *Yucca* species found there are both affected. However, in the case of *Y. baccata*, it is sympatric with other *Yucca* species, including Soaptree Yucca (*Y. elata)*, which apparently did not show this anomaly (or it wasn’t sampled). While it remains possible that sampling deficiencies have limited detection of the spatio-temporal extent of anomalous blooms, absences are well documented across the range in other areas over both the anomalous and normal flowering period, and no other years show strong evidence of concerted anomalies as seen in the fall and winter 2018.

A key question is the cause of anomalous blooms, and it has been speculated that a much colder and wetter than usual fall in 2018 across portions of the desert Southwest of the United States may have triggered this event^7^, but this has yet to be tested rigorously. Climate factors have been previously hypothesized to affect phenology of *Yucca*. Smith and Ludwig (1976), for example, found that *Y. elata* populations at a site near the USDA Jornada Experimental Range in southern New Mexico formed stalks almost a month later in 1973 compared to 1972, and speculated that this delay was caused by a wetter, cooler spring. However, Ackerman (1980) claimed that in *Y. schidigera*, flowering may be driven more by photoperiod. Our results cast doubt on a purely photoperiod-driven phenological response, given differences in photoperiod in December when desert *Yucca* species were found in bloom in 2018, versus typical bloom timing in March and April. Both suggest the importance of other proximal climatic drivers. However, we forego a more thorough test of such climate drivers here, since doing so properly requires a much more thorough examination of overall phenology across well-studied *Yucca* species and multiple years of data. Particular attention to localized areas of high rainfall and unusual cold, especially in relation to spatially restricted anomalous flowering, is especially warranted.

A final question we address here is whether anomalous *Yucca* blooming meant that areas where those blooms occurred had normal flowering patterns in the normal blooming period. If so, then such anomalies may have strong fitness consequences, especially since flower production is expensive in *Yucca*, and often plants cannot flower each year due to tradeoffs between optimizing vegetative versus floral growth (Smith and Ludwig, 1976). Figure 4 provides clear evidence that those areas with anomalous blooms also had plants flowering at normal times. While it is unlikely that anomalous flowers are pollinated given the presumed absence of its pollinator, this too requires a more thorough examination to verify. It may be that climatic cues are synchronized between *Yucca* and their obligate pollinator moth and that the unusual flowering timing is adaptive, allowing yuccas to take advantage of the right conditions for pollination. However, Rafferty et al. (2015) suggest that mutualisms between *Yucca* and their pollinating moths are not necessarily synchronized to climate cues, at least in normal spring flowering conditions. Further examination of whether any plants that were photographed formed fruits during the period between unusual and usual flowering would help provide evidence for the intriguing question of whether adult pollinators were also present.

### Caveats and Conclusions

Our work demonstrates that iNaturalist records provide a critical resource for phenology studies, if these are scored carefully following a well designed rubric. Here we have focused on the ability of iNaturalist records to uncover spatial and temporal trends in flowering, especially the ability to localize where and when anomalous *Yucca* flowering occurred in fall and winter 2018 after reports of such events in the media. While our results strongly show the value of iNaturalist data in answering such spatiotemporal phenology questions, we close with a few needed caveats regarding use of these data. First, while identifications are generally good for the focal taxa used in this study, despite the general challenge of field identification of *Yucca* (McKelvey and Sax, 1933), there are still cases where “research grade” specimens are misidentified. While evaluating identifications was not explicitly part of our efforts, and none of the participants in this study are experts in taxonomic identification of *Yucca* from photographs, we noted a very small percentage of obvious misidentification for the six focal species (well below 1 %). This low rate likely reflects in part our choice of particularly easily identified taxa such as *Y. brevifolia*. Still, as the *Y. filamentosa* panel in Figure 4 shows, there remain spatial outliers in iNaturalist that fall outside the known species range as documented by Flora of North America (2008), which likely indicate cultivated observations (discussed above) but not labelled as such, or potentially misidentifications. Finally, while extremely rare since most uploads come from cameras with automated date stamps, we did find cases (e.g. upload dates preceded dates of the photograph being taken) where camera date and times may be improperly set or dates are improperly entered manually.

Like all occurrence datasets reused from aggregators, care must be taken when using such data, in order to flag problem records and outlier information. Our efforts at vetting data and making our results immediately available on plantphenology.org provides not only a mechanism for re-use but also a means to assure that data can ultimately be improved. Finally, publishing iNaturalist records to plantphenology.org also extends data integration mechanisms across multiple types of data sources, which now include phenology observations from monitoring networks such as NPN, phenology annotations from herbarium specimens, and now citizen science-based photographic evidence from the iNaturalist platform. We also note the potential for such datasets to serve as inputs into rapidly developing supervised machine learning approaches (Lorieul et al. 2019) to scale up phenology reporting in the future.

## Acknowledgements

We thank the many citizen scientists using iNaturalist to photograph and identify occurrences of *Yucca*. Their efforts are the reason this work was possible. We also thank John Deck, who assisted with ingestion of the dataset from this effort into the Global Plant Phenology Data Portal (http://www.plantphenology.org/). We appreciate informal reviews of this work by Lucas Majure, Carrie Seltzer, and Shawn Taylor.

## Supplements

Supplement 1. A scoring guide for documenting *Yucca* phenological traits examined here: whole plant presence, flower presence and open flower presence.

## Footnotes

1 http://www.inaturalist.org/

2 https://www.desertsun.com/story/desert-magazine/2019/01/30/earlv-bloom-of-joshua-treescould-be-dire-for-mojave-desert-ecosvstem/2706708002/

3 https://www.usanpn.org/usa-national-phenoloqv-network

4 http://www.neonscience.org

5 http://www.plantphenology.org

6 https://github.com/biocodellc/ppo-data-pipeline

7 http://www.hidesertstar.com/news/article_18205bf4-f740-11e8-9c49-6fef1825a383.html

