## Supplement 1 for "Methods for broad-scale plant phenology assessments using citizen scientists’ photographs"

### ***Introduction***

Below we provide details about how Yucca iNaturalist records were scored, focusing on the key traits that allowed us to document flower and open flower presence and absence. Those traits are “whole plant presence”, “flower presence” and “open flower presence”.

All images used in this tutorial have a CC-BY copyright and we provide attribution by adding the URL in order to link back to the original record on iNaturalist.

### ***Whole Plant vs. Part of Plant***

For ground species, whole plant presence was based on if the middle of the plant where the flowering stalk grows from is visible. If there was no evidence of a living stalk out of frame, it was scored as a whole plant. If a dead stalk was present but out of the frame, it was also scored as a whole plant.

For branched, tree-like species, whole plant presence was based on if the tips of all the branches could be seen. In cases where branches clearly extend past the frame of the photograph, those were scored as “whole plant absent”.

The below images show examples of unambiguous whole plant presence and more challenging examples to score.

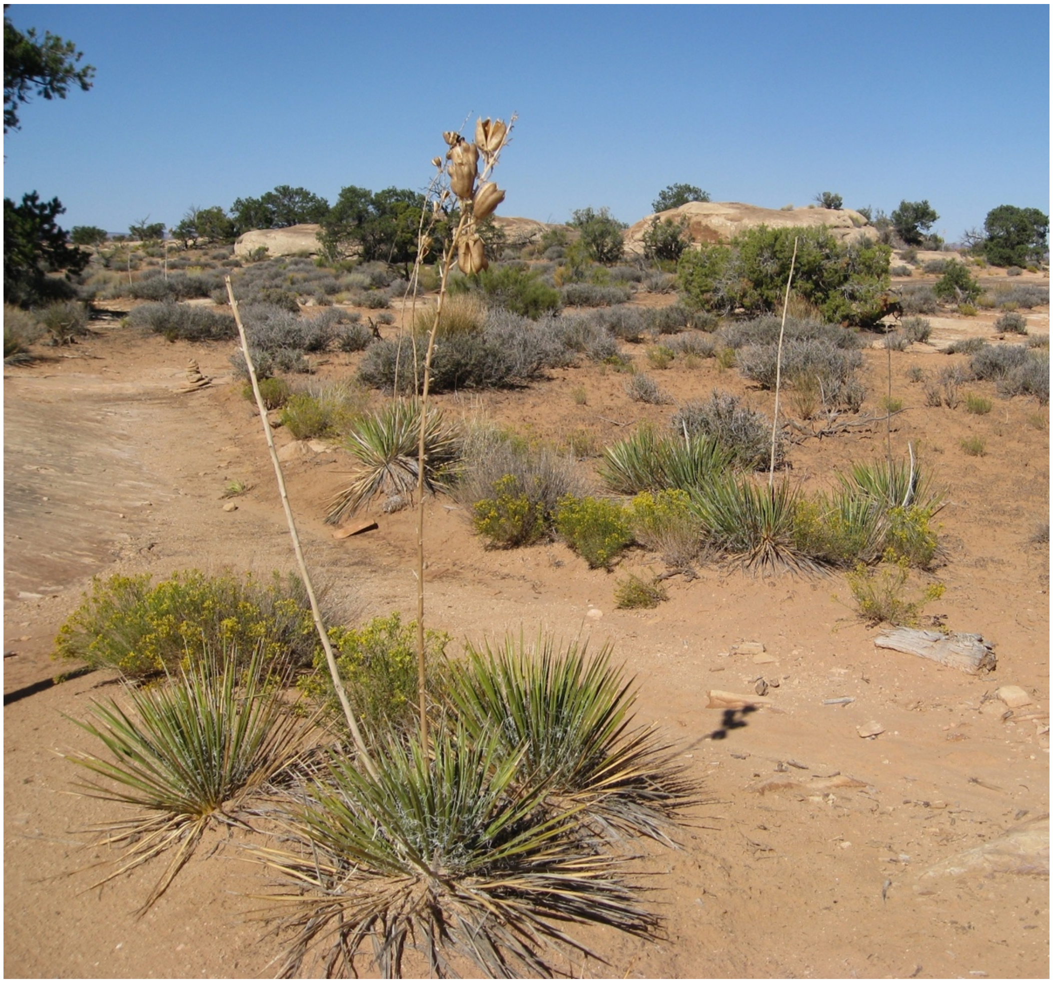

This *Y. angustissima* image shows an example of **whole plant present**.
Source: https://www.inaturalist.org/observations/117705

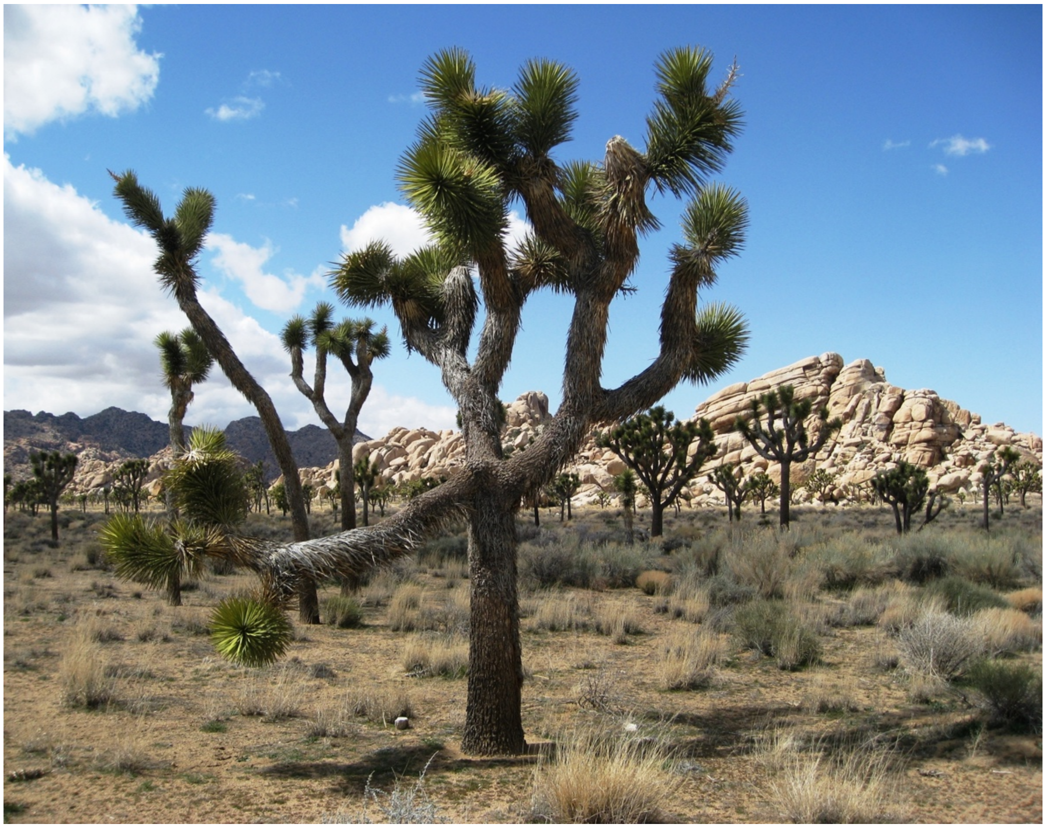

This *Y. brevifolia* image shows an example of **whole plant present** for an arborescent species.
Source: https://www.inaturalist.org/observations/113224

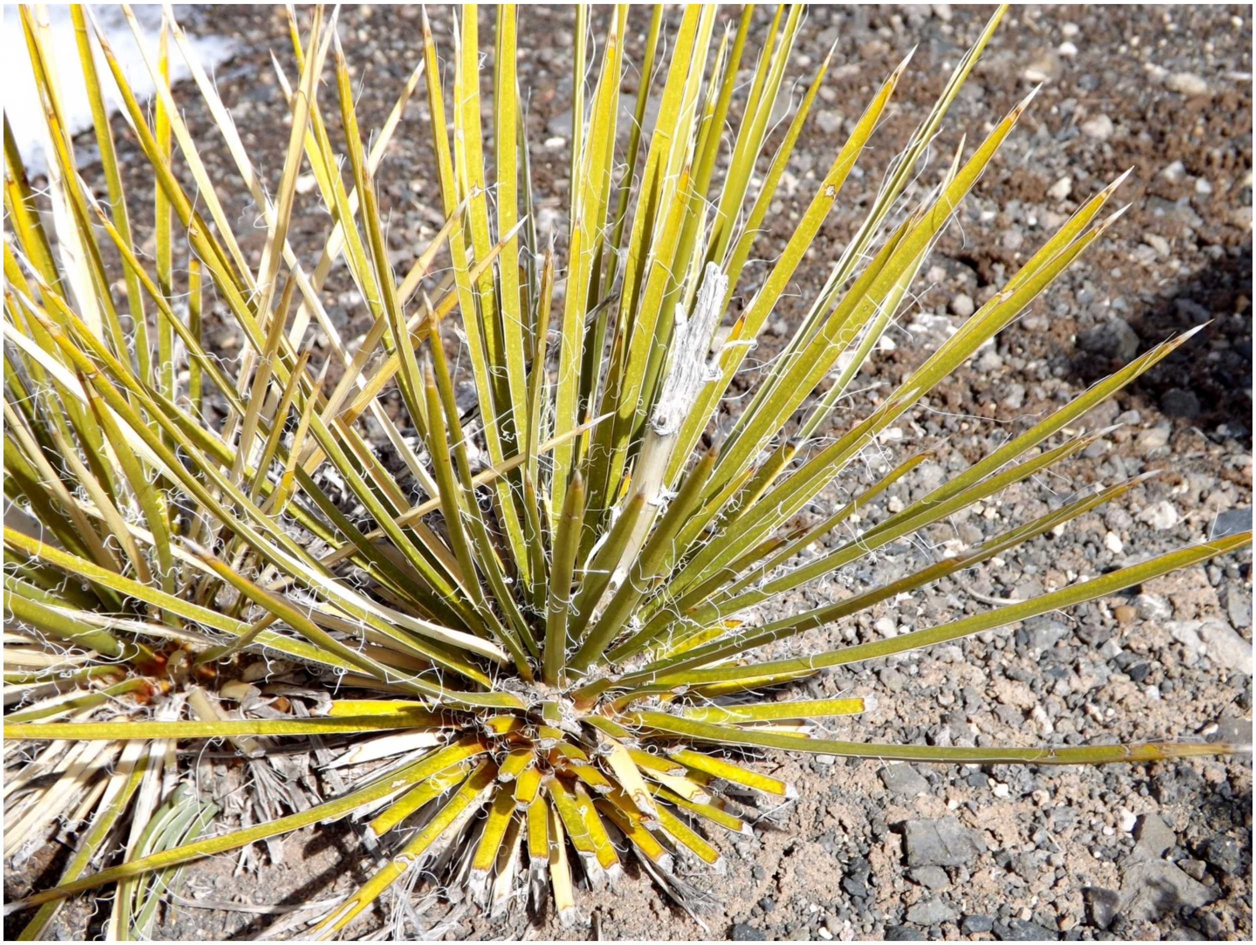

Though this image does not capture all of the leaves on this plant, this is another example of **whole plant present** because the bases of the leaves in the middle of the plant and no living stalk is out of the frame.
Source: https://www.inaturalist.org/observations/496349

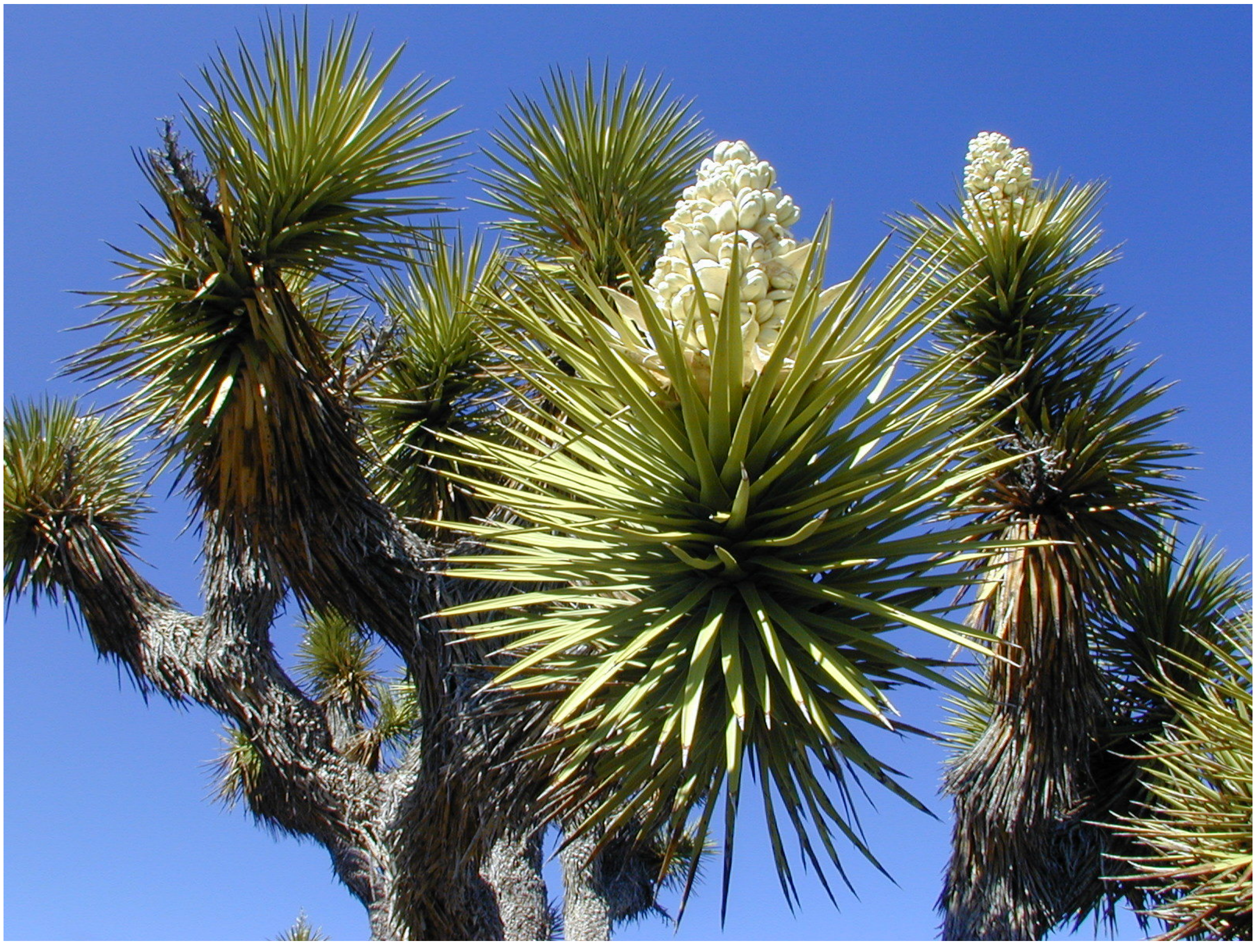

This *Y. brevifolia* image is an example of **part of plant**, especially given that we know *Y. brevifolia* has a treelike habit and all of the tips of the branches are not visible.
Source: https://www.inaturalist.org/observations/551671

### ***Flowers Present vs. Flowers Absent***

Flowers were considered to be present if at least one bud or non-senesced flower was visible on the target plant. Visible bracts do not mean a flower or bud is present. Examples are shown below.

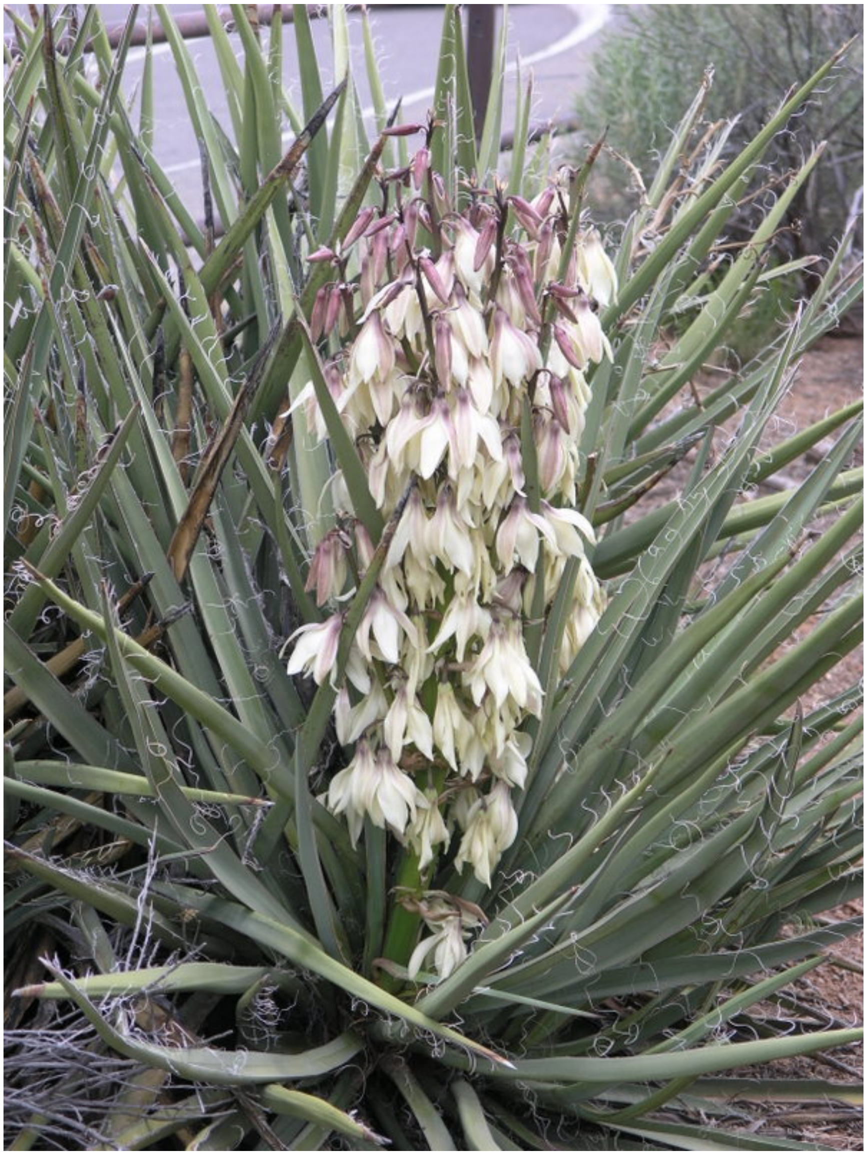

This image of *Y. baccata* has **open flowers present**, and therefore, also **flowers present**.
Source: https://www.inaturalist.org/observations/60429

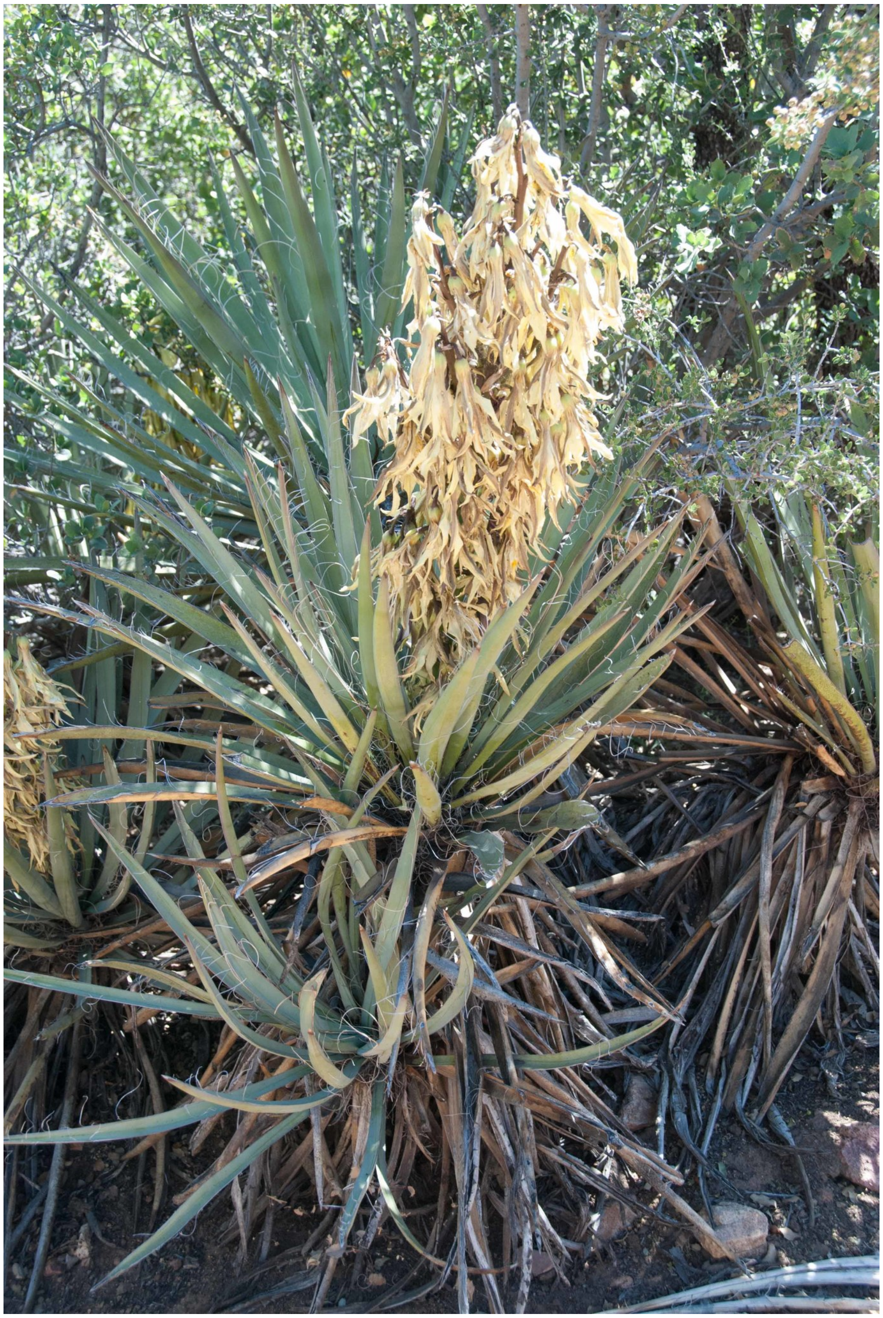

This image was scored as (non-senesced) **flowers absent**. We do not include senesced flowers, visible in the above photograph, in our scoring rubric as evidence of functional flowers present because they are not reproductively active.
Source: https://www.inaturalist.org/observations/6360640

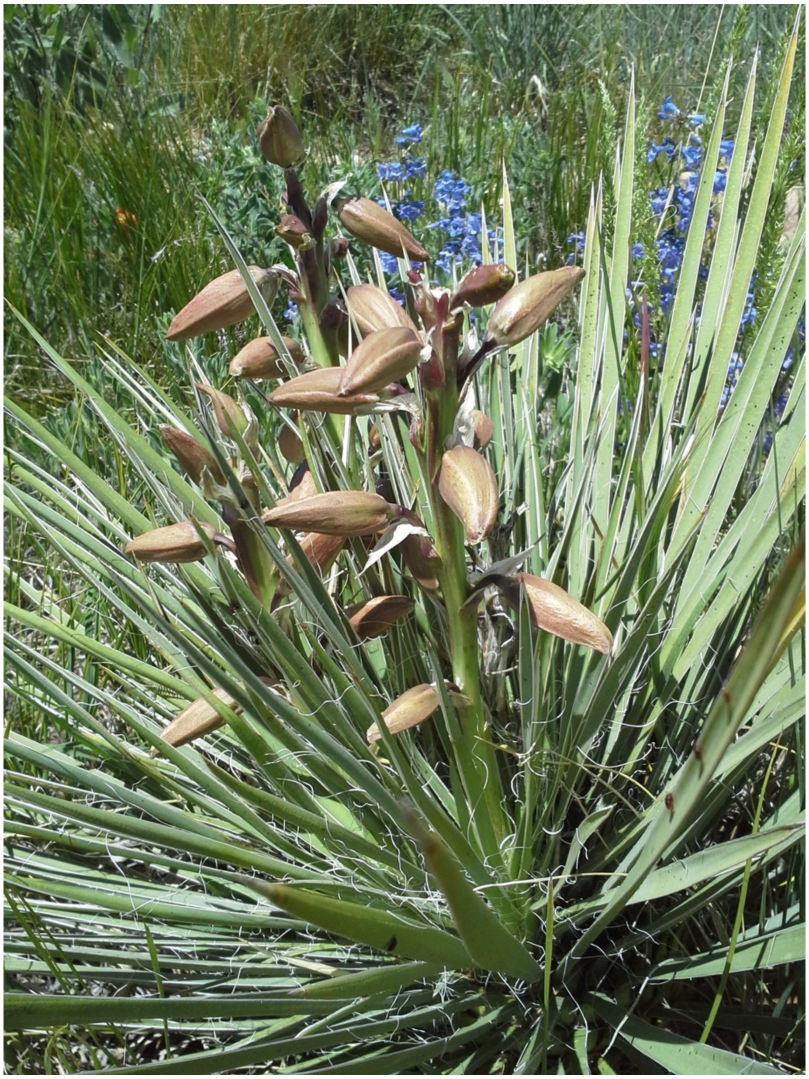

Flower buds here are clearly visible emerging from the bracts and therefore, this photograph was scored as **flowers present**.  However, open flowers are absent.
Source: https://www.inaturalist.org/observations/1056068

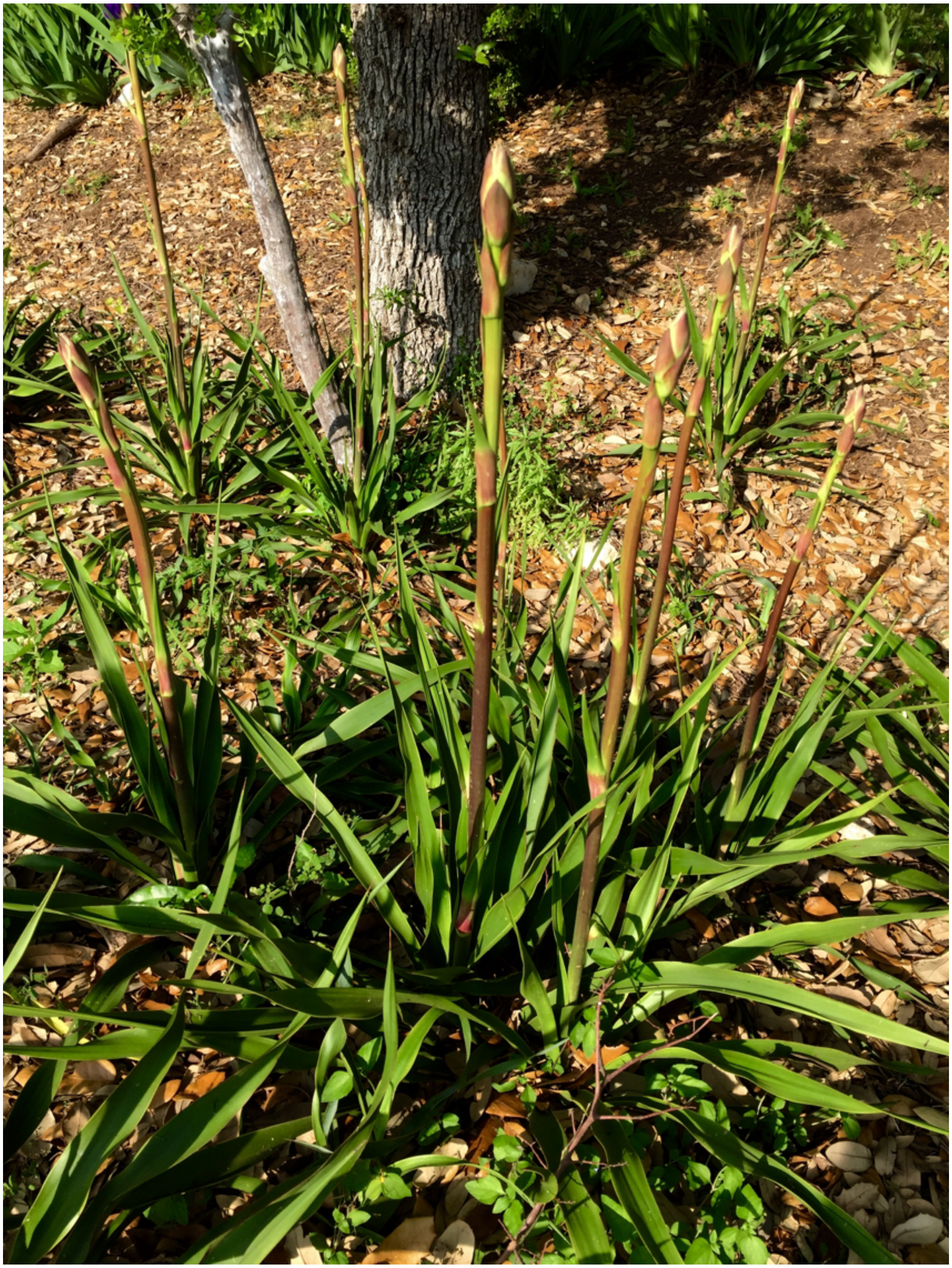

However, in this photograph, bracts are present but no buds are clearly visible, and so this is scored as **flowers absent.**
Source: https://www.inaturalist.org/observations/1362960

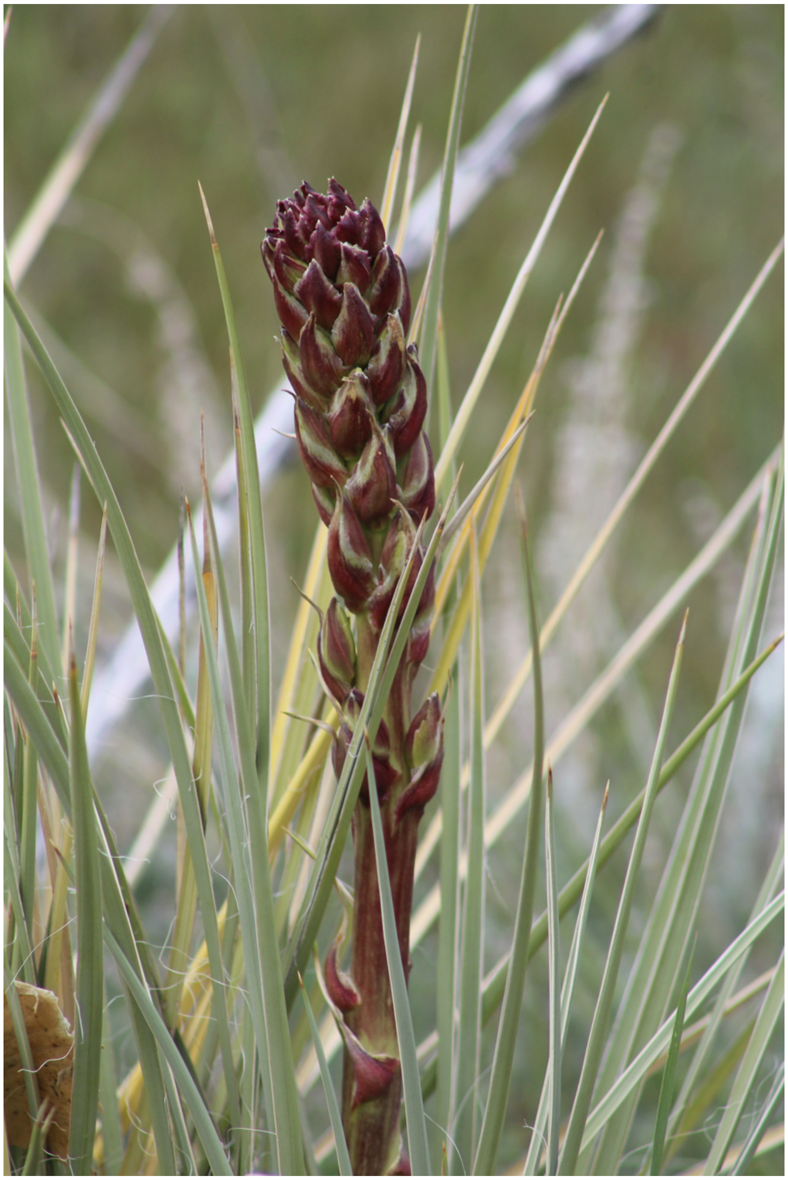

This is a more challenging case but this was scored as **flowers present**. The lower portion of the flowering stalk clearly has visible buds; the upper portion does not, but any visible buds is enough to score present.
Source: https://www.inaturalist.org/observations/26406605

***Open Flowers Present vs. Open Flowers Absent***

Open flowers were considered to be present if at least one open, non-senesced flower was visible in the photograph. Open flowers are defined as flowers where the reproductive parts or open flower parts such as petals are visible. Wilting, dried and discolored flowers were considered to be senesced. Lack of wilting and discoloration indicative of old flowers were key for determining non-senesced from senesced flowers.

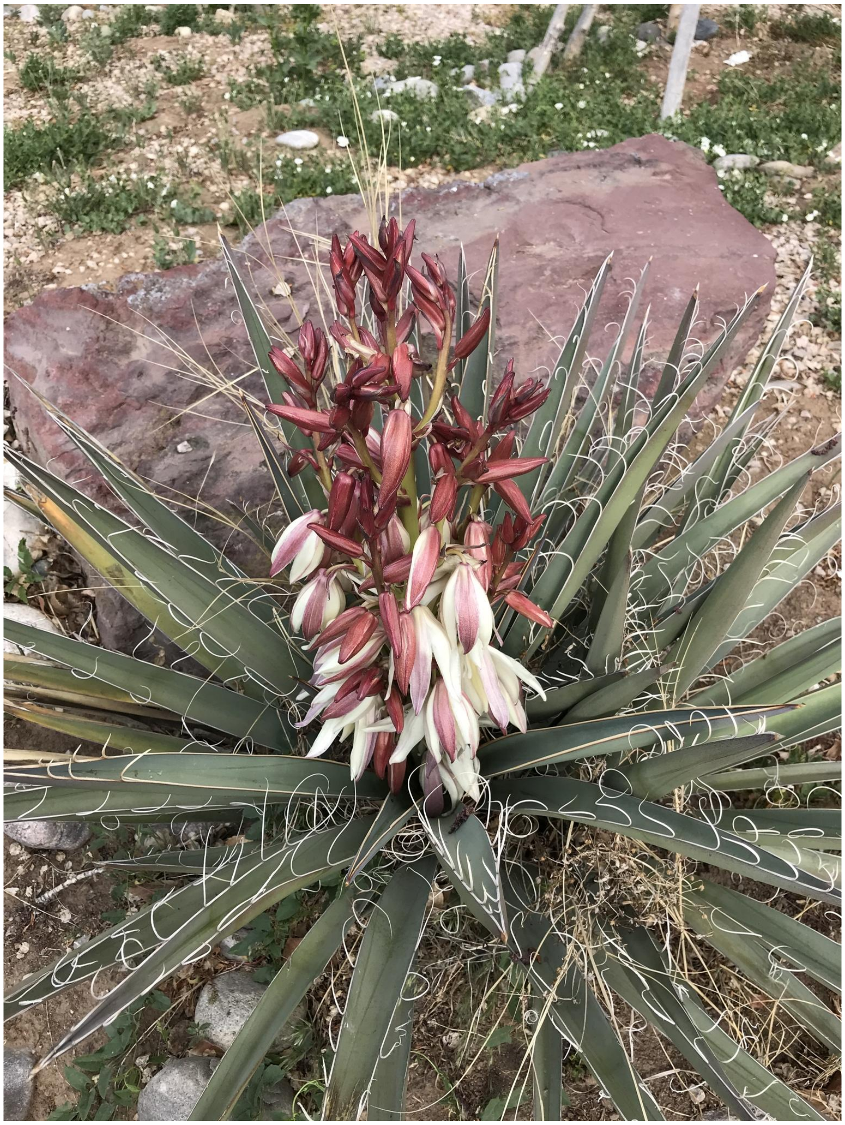

This image of *Y. baccata* clearly shows **open flowers present**. Though there are unopened buds on this plant, only one visible flower must be open for an image to satisfy the requirements for open flowers present. Note that while most yucca have white flowers, some species have dark-colored sepals. Scorers were only asked if open flowers are present on images scored as flowers present.
Source: https://www.inaturalist.org/observations/12725125

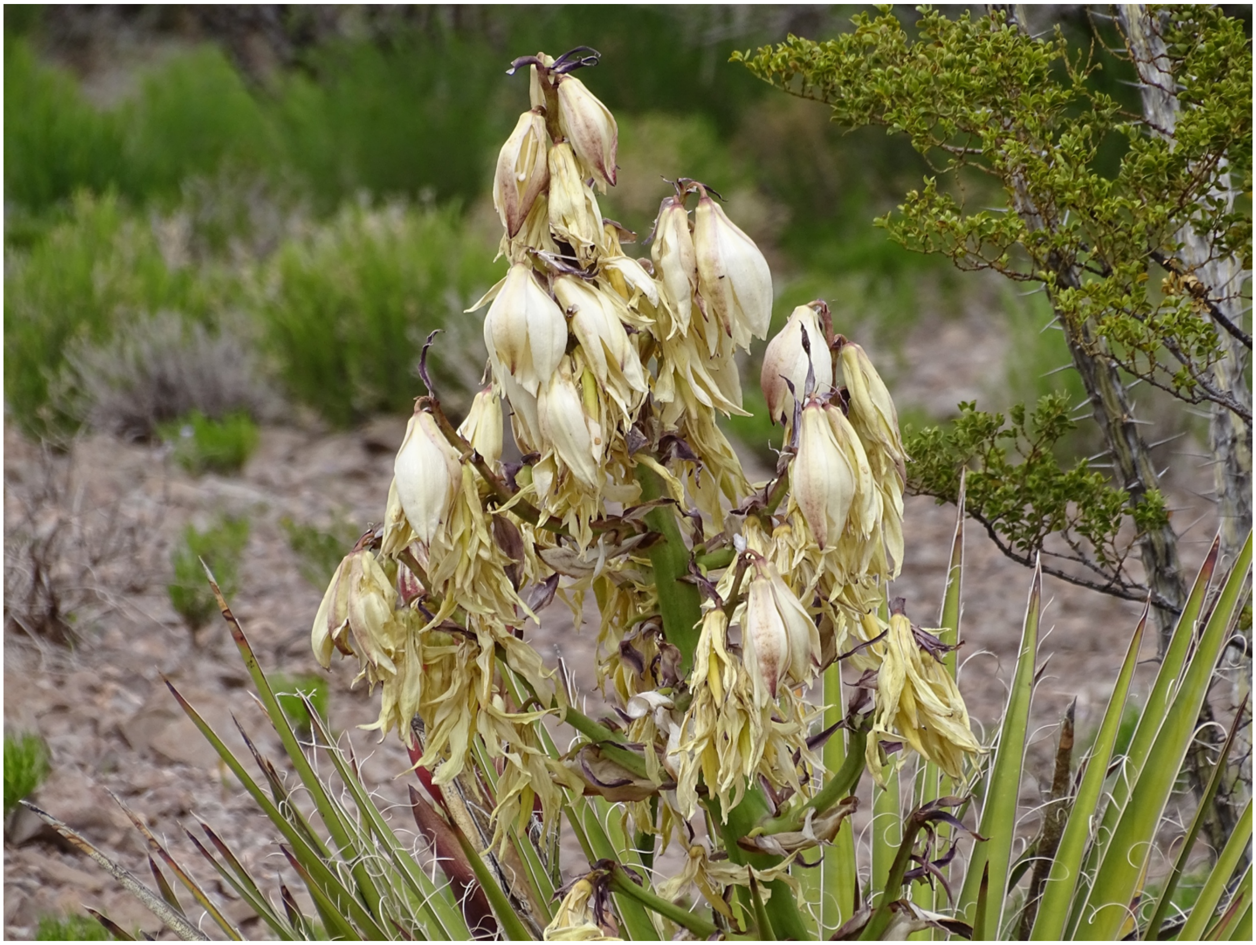

This image also has **open flowers present**. While there are senesced flowers as well, any clearly non-senesced flower, such as those near the top of the flower stalk, are enough to score as flowers present.
Source: https://www.inaturalist.org/observations/22398424

### ***Miscellaneous***

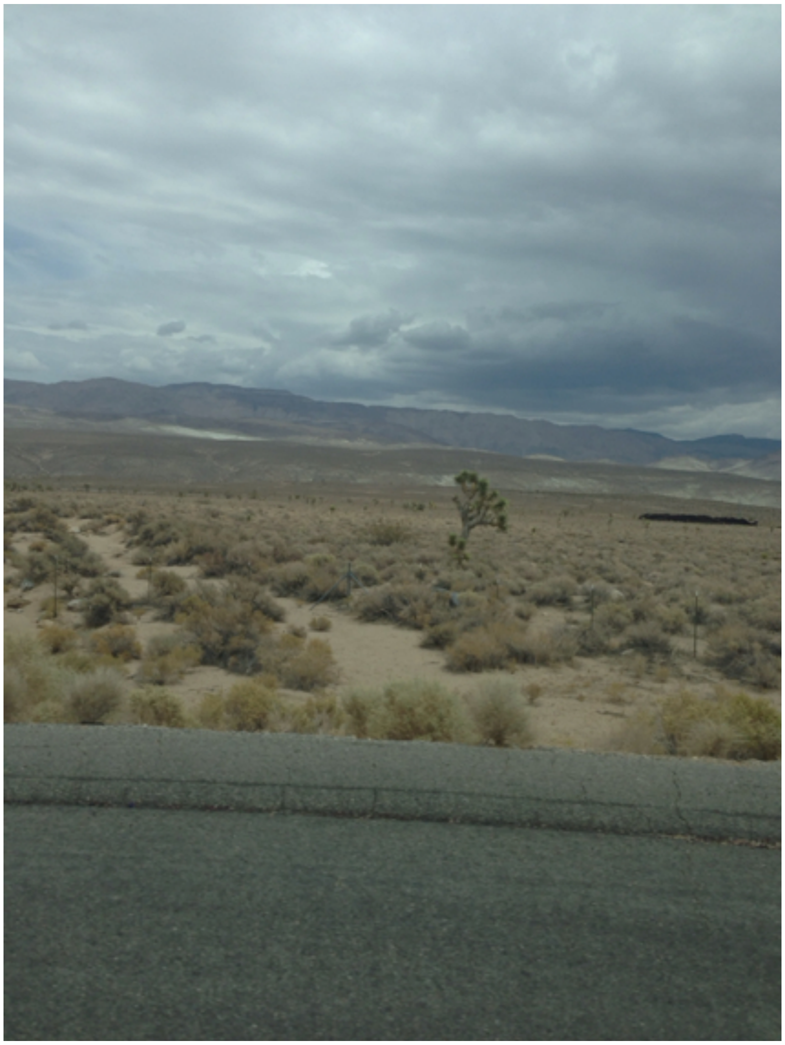

Finally, we included this image as an example for what was scored as “uncertain”. There are whole plants of (what is almost certainly) *Y. brevifolia* visible in this image, but the blurriness of the photo and distance from the plant makes determining the flowering phenology impossible. Source: https://www.inaturalist.org/observations/806851
